# The oriental fruit moth genome provides insights into evolutionary adaptation of tortricid pests

**DOI:** 10.1101/2020.09.04.282533

**Authors:** Wei Song, Li-Jun Cao, Lei Yue, Shao-Kun Guo, Jin-Cui Chen, Ya-Jun Gong, Xu-Lei Fan, Shu-Jun Wei

## Abstract

Moths of the family Tortricidae (Insecta: Lepidoptera) usually distribute in temperate and tropical high upland regions. The oriental fruit moth (OFM) *Grapholita molesta* is a globally important pest of stone and pome fruit from this family. In this study, we assembled a chromosome-level genome for the OFM and conducted a comparative genomic analysis with other lepidopterans. This genome was assembled to 28 chromosomes with a size of 517.71 Mb, an N50 of 19.5 Mb, and a BUSCO completeness of 97.1%. In total, 19968 protein-coding genes were predicted, among which 15269 were functionally annotated. We manually annotated ten gene families for 13 representative genomes from the Lepidopteran. In general, two tortricid moths have a moderate number of detoxification and receptor genes and the lowest number of HSP genes, congruent with their polyphagous dietary and pattern of distribution. However, the OFM has the highest number of P450, I Rs and ORs among all species. Compared to the other tortricid species of the codling moth *Cydia pomonella*, the OFM has more genes in all gene families. Our results indicate that the high number of some detoxification and receptor genes may be related to the strong adaptation of OFM as a global pest. The high-quality genome of OFM provides an invaluable resource for understanding the ecology, genetics, and evolution of tortricid moths.

## Introduction

The Tortricidae is one of the largest families of Lepidoptera with over 5000 described species, including many economically important pests in agriculture and forestry (van der Geest & Evenhuis, 1991). Tortricid moths cause damages to their hosts in larval stages by feeding on the leaves, fruits, buds, shoots, twigs, and stems. Moths of the family Tortricidae tend to distribute in temperate and tropical high upland regions (van der Geest & Evenhuis, 1991). However, some species expanded their distribution range as global invasive species (Kirk et al., 2013b; Wan et al., 2019).

The oriental fruit moth (OFM) *Grapholita molesta* is a cosmopolitan pest of stone and pome fruits from the Tortricidae (Neven et al., 2018; Quaintance & Wood, 1916; Torriani et al., 2010). Larvae of the OFM cause damage by boring into twigs as well as fruits (Figure 1). Population genetic analysis shows that this moth is native to East Asia and has spread into other stone-fruit growing continents in the past century (Kirk et al., 2013a; Song et al., 2018; Wei et al., 2015). Although many methods have been applied to counter outbreaks and spread of OFM, this remains a challenging work for both of the farmers and researchers (Audemard et al., 1989; Il’ichev et al., 2007; Rica et al., 2006; Rothschild, 1979; Sarker & Lim, 2018). Understanding evolutionary adaption underlying the worldwide distribution may provide insights into the management of this pest. Current studies identified pheromone gland gene, receptor genes, thioredoxin genes, detoxification genes, microRNAs from the OFM from transcriptome data to explore the possible mechanisms related to pest control (Chen et al., 2019a; Guo et al., 2017; Shen et al., 2018; Vatanparast & Kim, 2019; Wang et al., 2017). However, due to the lack of genomic resources, the molecular and genetic study on the OFM was largely delayed. Song et al. (2017) assembled a draft genome for the OFM based on sequences generated by the Illumina Miseq platform. However, because of the high fragmentation of this draft assembly, it was only applied for the development of microsatellite markers (Song et al., 2018; Song et al., 2017). A high-quality genome is essential to understand the evolutionary adaptation and develop new control approaches of OFM.

**Fig. 1.**
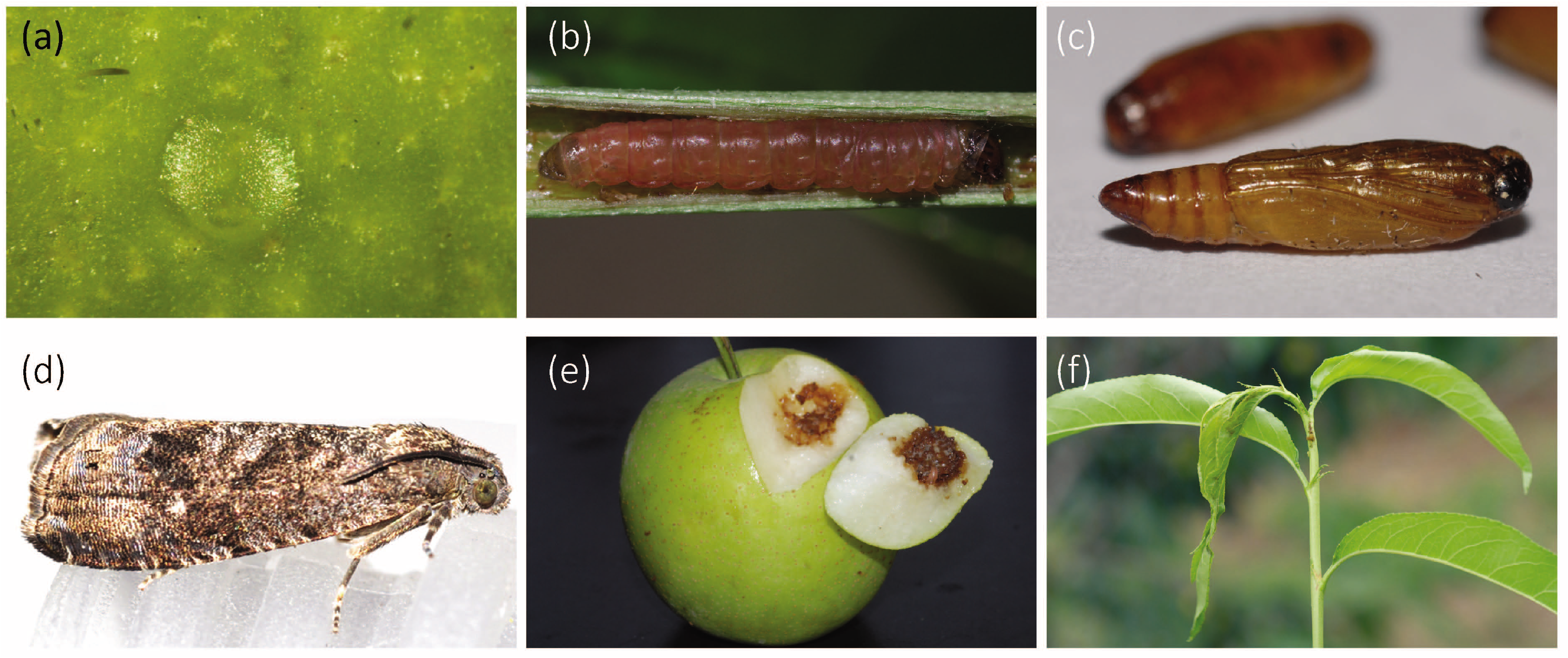
Image of four stages of *Grapholita molesta*. Egg (a), larva (b), pupa (c), adult (d), the damage symptoms on the fruit of pear (e) and the twigs of peach (f) were shown. Photographs were taken by Shu-Jun Wei.

In the study, we report a chromosome-level genome of OFM. This genome was de novo assembled based on sequences obtained from the PacBio and Illumina platforms and assembled at the chromosome level assisted by the Hi-C technique. We compared genome features between OFM and another cosmopolitan pest from Tortricidae, the codling moth *Cydia pomonella* (Lepidoptera: Tortricidae) (Wan et al., 2019), to explore the evolutionary adaptation of tortricid moths. The genomic resource developed here provides an invaluable resource for understanding the genetics, ecology, and evolution of moths.

## Materials and methods

### Sample preparation, DNA libraries construction and sequencing

Laboratory reared strains of the OFM were used for de novo genome sequencing. The OFM strain derived from three pairs of males and females has been maintained for about 10 generations on apples in the Beijing Academy of Agriculture and Forestry Sciences. Genomic DNA was extracted using the DNeasy tissue kit (Qiagen, Hilden, Germany) for Illumina libraries and using the MagAttract HMW DNA kit (Qiagen, Hilden, Germany) for NanoPore library construction. The Illumina libraries were sequenced using the Illumina HiSeq X Ten platform (Illumina, CA, USA) and the NanoPore library was sequenced using the Oxford NanoPore platform (Pacific Biosciences, CA, USA) (Table S1).

The Hi-C proximity ligation library was constructed following Berkum et al. (2010). The genome was digested by restriction enzyme Mbol. Fragments were sheared into 200-600 bp. The fragment ends were added A-tailing by Klenow (exo-) and then added Illumina paired-end sequencing adapter. The libraries were amplified by 12 cycles PCR and sequenced in the Illumina HiSeq X10 platform.

### Transcriptome sequence

In total, 4 RNA-seq libraries were constructed for eggs, larvae, pupae and adults of the OFM (Table S1). All libraries were prepared using VAHTSTM mRNA-seq V2 Library Prep Kit for Illumina according to the manufacturer’s instructions (Vazyme, Nanjing, China) and sequenced using the Illumina NovaSeq platform. The RNA-seq reads were de novo assembled and mapped to the OFM genome using TopHat (Trapnell et al., 2009) with default parameters. The assembly results of RNA-seq were used as evidence for genome structure annotation. After removing the low quality reads with Trimmomatic v0.38 (Bolger et al., 2014), the reads were mapped to the chromosome-level genome using Hisat v2.2.0 (Kim et al., 2019) and assembled with StringTie v2.1.2 (Pertea et al., 2015). FPKM (Fragments Per Kilobase per Million) values of each annotated gene in each RNA-seq were estimated with cufflinks v2.2.1 (Kim et al., 2013).

### Genome size estimation, assembly, and quality assessment

The raw reads generated from the Illumina platform were filtered by Trimmomatic v0.38 (Bolger et al., 2014). To estimate the genome size of the OFM, we used paired-end sequencing reads from Illumina sequencing library to determine the distribution of K-mers. Genome size, heterozygosity and rate of duplication were estimated using the online GenomeScope v1.0 software (Vurture et al., 2017).

Long reads generated from the NanoPore platform were corrected and assembled using CANU version v1.8 (Koren et al., 2017) with default parameters. The assembled contigs were polished based on Illumina short reads using Pilon v1.22 (Walker et al., 2014). To remove the possible secondary alleles, the assembled contigs were filtered by using the pipeline Purge Haplotigs (Roach et al., 2018), resulting in a contig-level genome. The Illumina short reads sequenced from the Hi-C library were used to assemble these contigs into a chromosome-level genome using the Juicer v1.5 (Neva C. et al., 2016) and 3D de novo assembly (3D-DNA) pipeline (Dudchenko et al., 2017) (Table 1).

**Table 1.** Completeness of major versions of assembly and annotation of *Grapholita molesta* genome estimated by BUSCO

| Step | Version | Software | Complete % | Single copy % | Duplication % | Fragment % | Missing % | Size (Mb) / No. gene | N50 (Mb) |
| --- | --- | --- | --- | --- | --- | --- | --- | --- | --- |
| Assembly | Contig assembly | CANU | 83.3 | 57.7 | 25.6 | 7.1 | 9.6 | 949.05 | 0.85 |
|  | Syteny reduction | purge_haplogs | 76.6 | 74.7 | 1.9 | 10.5 | 12.9 | 521.60 | 1.79 |
|  | Contig polish | PILON | 96.7 | 93.1 | 3.6 | 1.1 | 2.2 | 524.93 | 1.79 |
|  | Linkage group assembly | Juicer + 3D-DNA | 93.3 | 92.2 | 1.1 | 1.0 | 5.7 | 525.22 | 19.5 |
|  | Linkage group polish | PILON | 97.1 | 96.1 | 1.0 | 1.0 | 1.9 | 517.71 | 19.5 |
| Annotation | Annotation | MAKER | 90.1 | 86.1 | 4.0 | 6.6 | 3.3 | 39,274 | NA |
|  | Filtering based on expression | In-home script | 89.8 | 85.9 | 3.9 | 6.1 | 4.1 | 19,478 | NA |
|  | Gene structure improvement | PASA | 92.6 | 76.6 | 16.0 | 4.4 | 3.0 | 19,968 | NA |
|  | Functional annotation | EGGNOG | 92.6 | 76.6 | 16.0 | 4.3 | 3.1 | 15,269 | NA |

Note: X/Y in Genome size (Mb) / Gene number items represent total size/N50 length. NA, not available.

Completeness of each assembled version of the genome was assessed using a Benchmarking Universal Single-Copy Orthologs (BUSCO) v3.0.2 (Simao et al., 2015) analysis, based on the insecta_odb9 database (1,658 genes). We conducted a synteny analysis between OFM and another tortricid moth, the codling moth *Cydia pomonella* (Assembly accession: GCA_003425675.2) (Wan et al., 2019) and *Spodoptera litura* (Assembly accession: GCF_002706865.1) (Cheng et al., 2017) using TBtools v0.58 (Chen et al., 2020) to check the quality of the chromosome-level genome. Short contigs in *S. litura* and *C. pomonella* genomes were removed before the blast alignment.

### Repeat elements and non-coding RNAs annotation

Repeats and transposable element families in the OFM genome were first detected by RepeatMasker pipeline v4.0.7 (Tarailo-Graovac & Chen, 2009) against the Insecta repeats within RepBase Update (http://www.girinst.org) and Dfam database (20170127), with RMBlast v2.10.0 as a search engine, which employs complementary computational methods to build and classify consensus models of putative repeats. tRNAs were annotated by tRNAscan-SE (Lowe & Eddy, 1997) with default parameters. rRNAs were annotated by RNAmmer prediction (Lagesen et al., 2007).

### Protein-coding gene annotation

The protein-coding gene in the OFM genome was structurally annotated using *ab initio*, RNA-seq-based, and homolog-based methods. First, the RNA-seq reads were aligned to the reference genome using PASA (Brian J et al., 2003) to extract the complete genes for SNAP model training using SNAP v2013-02-16 (Korf, 2004). Second, a combination of de novo prediction using SNAP and Augustus v3.2.3 (Stanke & Waack, 2003), homolog-based prediction using protein-coding genes of *Drosophila melanogaster* and *Bombyx mori*, and RNA-seq-based prediction using transcripts of OFM was conducted using Maker v3.01.03 genome annotation pipeline (Cantarel et al., 2008). For Augustus analysis, the gene model from BUSCO analysis of chromosome-level assembly was used. Third, the gene set annotated by the Maker pipeline was filtered by keeping those with an FPKM (fragments per kilobase of exon model per million reads mapped) value > 0 in any RNA-seq library. Finally, PASA was used to update the annotation results with transcripts of OFM; all predictions were further filtered using GffRead v0.11.7implemented in Cufflinks v2.2.1 (Kim et al., 2013) to remove without or with multiple stop codons.

Functions of the protein-coding genes were annotated using eggNOG-Mapper v1.0.3 (Huerta-Cepas et al., 2017) against the database EggNOG v5.0 (Huerta-Cepas et al., 2019). Genes that can be functionally annotated by EGGNOG analysis were retained in gene structure annotation and used for further analysis.

### Orthology identification and gene family expansion/contraction analysis

Protein-coding genes from available genomes of two Coleoptera, two Diptera and other 11 Lepidoptera were retrieved from the NCBI genomes database and published references for comparative analysis. Orthologs were identified using OrthoFinder v2.2.7 (Emms & Kelly, 2015) under default parameters. MAFFT v7.450 (Katoh & Standley, 2013) was employed to align amino acid sequences of 1:1:1 orthologous gene with the G-INS-I algorithm. The phylogenetic tree was inferred using an approximately-maximum-likelihood method implemented in FastTree v2.1.10 (Price et al., 2009), which is a default phylogenetic inference method for genomic data in OrthoFinder. We used r8s (Sanderson, 2003) to estimate the divergent times among species with divergence times of two nodes, i.e. *Tribolium castaneum* and *Anoplophora glabripennis* (Wang et al., 2019), *Trichoplusia ni* and S. *litura* (Wan et al., 2019) as calibrations. The Computational Analysis of gene Family Evolution (CAFE) version 3.1 (Bie et al., 2006) was used to analyze gene family expansion and contraction. The orthologue groups that present or absent in only one organism were defined as species-specific or lost orthologues. The gene sets groups with multiple-copy displayed in one organism were defined as specific expansions.

### Gene family annotation

Adaption to heat stresses usually relies on Heat shock proteins (HSP) (Bedulina et al., 2013; Colinet et al., 2013), while host plant adaptation usually employs detoxification genes to detoxify plant secondary metabolites in insects. There are five major detoxification gene families in animals, i.e., cytochrome P450 monooxygenases (P450s), glutathione-s transferases (GSTs), ATP-binding cassette transporters (ABCs), UDP-glycosyltransferases (UGTs) and carboxyl/cholinesterases (CCEs). Chemical senses to olfaction and taste have been found to play important roles in insects to find mates, food, and oviposition sites, and to avoid harmful situations and non-host habitats (Hansson & Stensmyr, 2011). Four gene families are often involved in chemical senses, including olfactory receptor (ORs), gustatory receptor (GRs), Ionotropic receptors (IRs) and Odorant-binding proteins (OBPs). Here, we manually annotated and compared those gene families among 13 representative genomes of Lepidoptera, to explore the possible genomic signatures related to the environmental adaption of the OFM as well as the other tortricid moth, *C. pomonella*.

We used both model-based and similarity-based methods to annotate the gene families. For model-based identification, the Hidden Markov models (HMMs) were downloaded from Pfam 32.0 database (September 2018; (El-Gebali et al., 2018)) and run with HMMER v3.3 (Finn et al., 2011). The corresponding HMM model not found in Pfam database was manually trained using HMMER under the default parameters. For similarity-based identification, orthologs from *D. melanogaster, B. mori, Aedes aegypti, Anopheles gambiae* and *C. pomonella* search against target genomes using BLAST v2.2.31 (Altschul et al., 1990) with an e-value cutoff of 1e^-5^. An automatic BITACORA v1.0 (Vizueta et al., 2020) pipeline (full mode) was used to conduct the HMMER and BLAST analyses. The annotated genes were filtered manually based on gene length and the presence of conserved domains by removing genes shorter than 80 amino acids and those lack of conserved domains.

## Results and discussion

### Genome sequencing and assembly

By integrating both Illumina HiSeq short-read sequencing and NanoPore long-read sequencing, we generated 102.43 Gb of clean read data, representing 197.85-fold coverage of the assembled 517 Mb genome (Table S1).

Based on Illumina short sequencing reads, the estimated size of OFM genome ranges from 393 Mb to 439 Mb; heterozygosity ranges from 0.721% to 0.865%; duplication rate ranges from 1.55% to 1.73% when K value was set to 17, 21, 27 and 31 (Figure S1). The heterozygosity of OFM is lower than the Thrips palmi (1.01% to 1.32%) (Guo et al., 2020) and the caddisfly *Stenopsyche tienmushanensis* (1.05%-1.10%) (Luo et al., 2018), higher than the beet armyworm, *Spodoptero exigua* (0.59%) (Zhang et al., 2019), and similar to the invading fall webworm (0.75% and 0.83%) (Wu et al., 2018).

The total length of the CANU assembly was 949.05 Mb, with an N50 equal to 850.95 Kb. After removing the possible secondary alleles, the size of the assembly was 521.6 Mb with an N50 length of 1.79Mb. Then, the contigs were polished into contig-level sequences using the short-reads again and generated a 524.93 Mb-length genome. These contigs were assembled into 28 chromosomes using Hi-C assembly, with a total length of 525.22 Mb and an N50 length of 19.5 Mb (Figure 2a). The chromosomal-level genome was finally polished by PILON to obtain a total length of 517.71 Mb and an N50 about 19.5 Mb (Table 1).

**Fig. 2.**
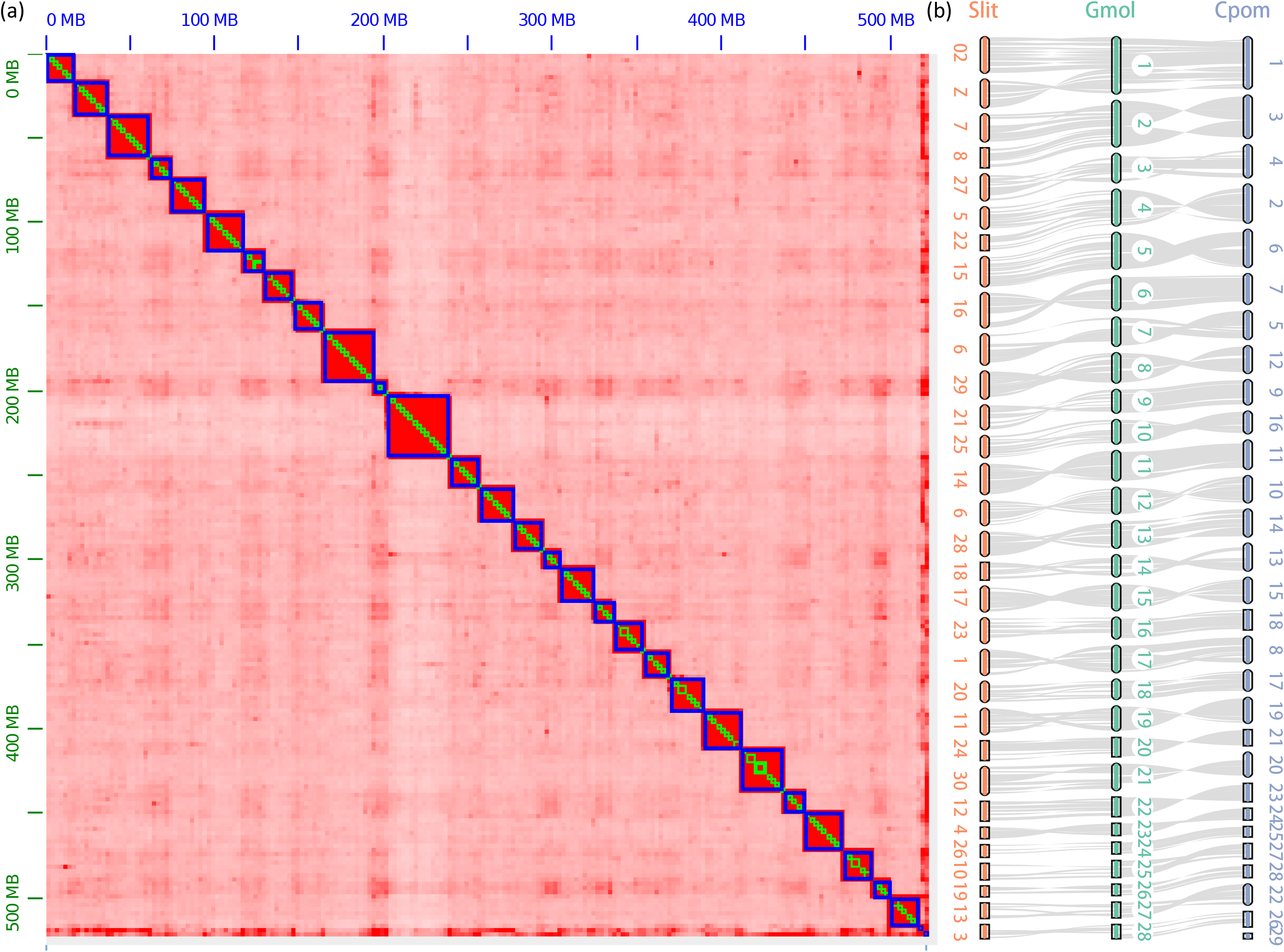
Genome-wide all-by-all Hi-C matrix of OFM (a) and Synteny blocks among *Grapholita molesta, Spodoptera litura* and *Cydia pomonella* genomes (b). The genome sequences were assembled using the Juicer grouped using 3D-DNA; in total 16 linkage groups were identified based on Hi-C contact, indicated by blue boxes. Only sequences anchored on chromosomes were shown in the plot. The orange, green and blue oval blocks represent chromosomes assembled in S. *litura, G. molesta* and *C. pomonella*; the gray lines represent the synteny blocks between the genomes. Powerful synteny blocks were identified between the three moths’ genomes.

To validate the quality of the assembled genome, we analyze the BUSCO completeness of each version of the OFM genome assembly. The completeness rises from 76.6 % to 97.1 %, with 96.1 %, 1.0 %, 1.0 %, and 1.9 % of genes were single-copy, duplicated, fragmented, and missing for the final genome assembly, respectively (Table 1). The quality of the assembly was much improved in each step, both in N50 size and BUSCO completeness. Our study showed that subsequent analyses of secondary allele removing and polish should be incorporated after assembly by traditional assembly methods.

This is the second chromosomal-level genome from the family Tortricidae following *C. pomonella* (Wan et al., 2019). Compared to the contig N50 of currently assembled genomes from the Lepidoptera (0.013 to 0.86 Mb) (Wan et al., 2019), the quality of contig level assembly is the highest in the OFM (1.79 Mb). The BUSCO completeness of chromosome-level assemblies for the lepidopteran species ranges from 91.5% to 98.5% (Wan et al., 2019), the completeness of OFM assembly is among the high-quality ones (97.1%). These results suggest that we have developed a high-efficiency genome assemble pipeline and the OFM genome assembly was of high quality in this study.

### Synteny and chromosome fusion in the Tortricidae

Powerful syntenic blocks were found between the OFM and *C. pomonella* genome as well as *S. litura* (Figure 2b). Every chromosome was one-to-one synteny between OFM and *C. pomonella* genome. In *C. pomonella* genome, there are 28 autosomes and one sex chromosome W. However, we have not identified the sex chromosome W (chrom 29 in *C. pomonella)* in this OFM genome. The number of chromosomes in the OFM is the same to *C. pomonella*. In Tortricidae, the number of chromosomes ranges from 27 to 31 (SuomMainen, 1969). The subfamily Tortricinae has 30 chromosomes, while the Olethreutinae, which includes the OFM and *Cydia pomonella*, has 28 chromosomes (van der Geest & Evenhuis, 1991). The results indicate we have successfully assembled all chromosomes except for the sex chromosome W in the OFM.

Fusion event between the sex chromosome Z and an autosome corresponding to chromosome 15 in the *Bombyx mori* reference genome was reported in the *C. pomonella* (Fuková et al., 2005; Wan et al., 2019). This phenomenon is also found in the OFM genome, confirmed the common fusion event in tortricid moths (Nguyen et al., 2013). Two additional fusion events were found in the OFM, as reported in the *C. pomonella* (Wan et al., 2019). The first one is the fusion of chromosome 7 and chromosome 8 of S. *litura* into Chromosome 2 of OFM; the second is the fusion of chromosome 5 and chromosome 22 of S. *litura* into chromosome 4 of OFM (Figure 2b).

### Protein-coding, non-coding RNA genes and repetitive elements

We annotated 39274 protein-coding genes using the MAKER pipeline, of which 19478 were supported by transcriptome data. We saved 19968 genes when the final PASA pipeline was finished, with the completeness of 92.6% estimated by BUSCO. Fifteen thousand two hundred sixty-nine gene sets were functionally identified in the genome of OFM using EGGNOG (Table 1).

We predicted 61 8s_rRNAs, 63 5s_rRNA and 29786 tRNAs (including 8853 tRNAs decoding standard 20 amino acid, 14 selenocysteine tRNAs (TCA), 13 possible suppressor tRNAs (CTA, TTA), 163 tRNAs with undetermined/unknown isotypes and 20743 predicted pseudogenes, in which 203 tRNAs with intron) in the OFM genome (Table S2).

A total of 15.37 Mb repetitive elements occupying 11.01% of the OFM genome were annotated using a combination of structural information and homology prediction (Table S3). Retrotransposable elements, known to be the dominant form of repeats, constituted a large part of the genome and included the most abundant subtypes, such as long terminal repeat elements (LTRs), long interspersed nuclear elements (LINEs) and short interspersed nuclear elements (SINEs). We predicted 41362559 bp retroelements (including 4947739 bp SINEs, 33924345 bp LINEs and 2490475 bp LTRs) in OFM genome based on Insecta RepBase libraries, which grouped in 231642 elements and occupied 7.99% of the assembly. We annotated 8670 DNA transposons (1730875 bp, 0.33 %), 19652 small RNA (4014531 bp, 0.78 %) and 135317 Tandem Repeats (TRs) (2 satellites and 135315 simple repeats). Nineteen thousand two hundred eighteen low complexity repeat elements occupying 0.18 % of the whole genome were identified (Table S3).

### Comparative genome analysis and phylogenetic relationships

A total of 320445 genes among 339228 genes from the 17 species were assigned to 14425 orthologue groups. We found 1785 genes in 427 orthologs were species-specific and there were 4948 orthologs present in all species. The G50 (The number of genes in the orthogroup such that 50% of genes are in orthogroups of that size or larger) and O50 (The smallest number of orthogroups such that 50% of genes are in orthogroups of that size or larger) of all genes was 28 and 3286 respectively (Table S4). For OFM, there were 19011 genes identified and 18258 genes occupying 96% were assigned in orthologs. A total of 117 gene families were species-specific and clustered into 13 orthogroups.

Phylogenetic analysis supports the monophyly of all involved superfamilies of Lepidopteran. Molecular dating support that OFM and *C. pomonella* diverged at 85.48 million years ago (Mya) (Figure 3). There were 883 gene family expansion/contractions in OFM genome, similar to *C. pomonella* (928) as well as most other moths and bufferflies (Table S5).

**Fig. 3.**
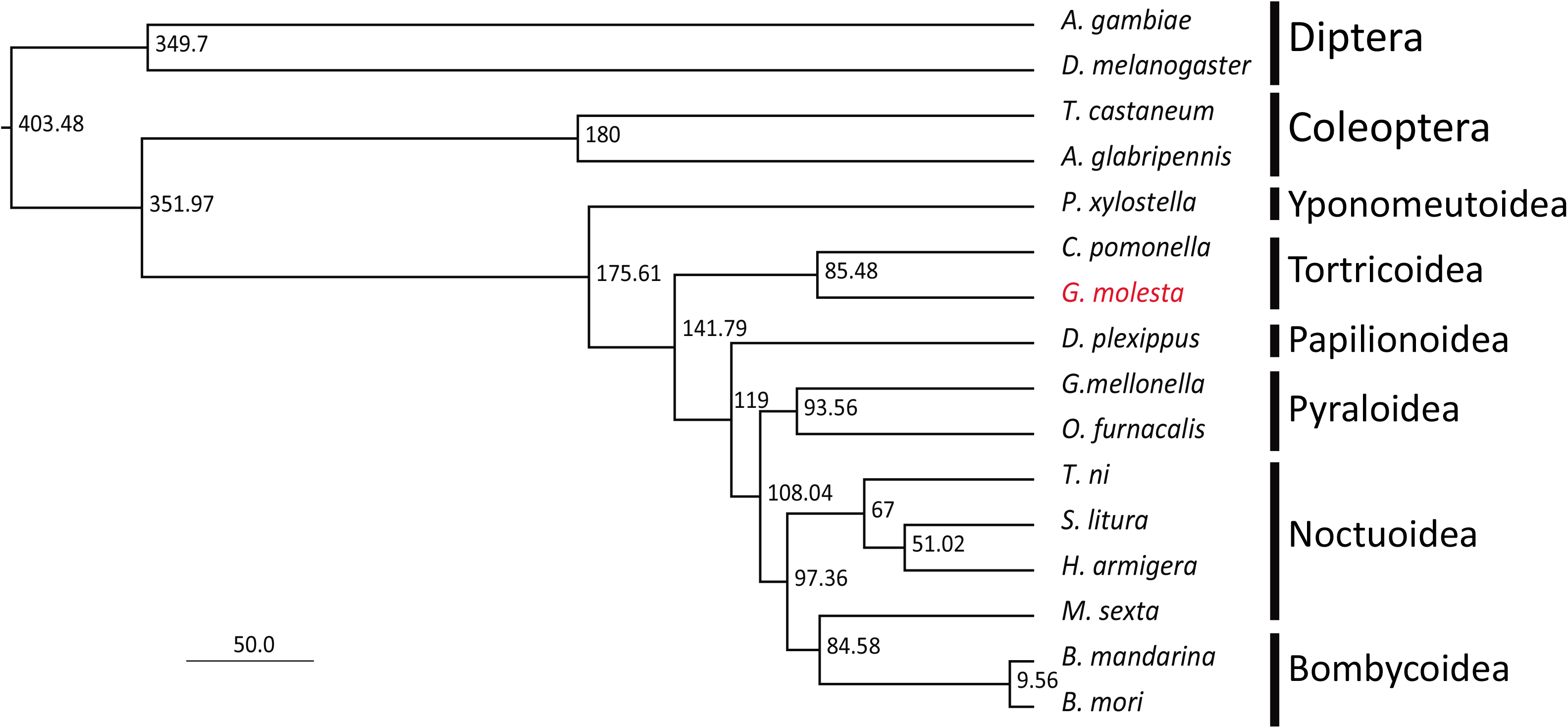
Phylogenetic relationships among *Grapholita molesta* and lepidopteran insects. Values on each node indicates the estimated divergence time. Superfamilies of 13 species of Lepidoptera and orders of four outgroups were indicated in the tree.

### Detoxification, receptor and HSP genes

In general, two tortricid moths, i.e. OFM and *C. pomonella* have moderate number of detoxification and receptor genes and the lowest number of HSP genes (Table 2), congruent with their polyphagous dietary and pattern of distribution (van der Geest & Evenhuis, 1991). The OFM has the highest number of P450, IRs and ORs among all species. Compared to the other tortricid species of the codling moth *Cydia pomonella*, the OFM has more genes in all gene families. Our results indicate that the high number of some detoxification and receptor genes may be related to the strong adaptation of OFM as a global pest.

**Table 2.** Number of genes in five detoxification families across 12 genomes from the Lepidoptera

| Species | P450 | GST | CCE | ABC | UGT | GR | IR | OBP | OR | HSP | Reference |
| --- | --- | --- | --- | --- | --- | --- | --- | --- | --- | --- | --- |
| Pxyl | 163 | 55 | 85 | 219 | 38 | 68 | 61 | 91 | 225 | 56 | You et al. (2013) |
| Gmol | 186 | 56 | 119 | 128 | 47 | 95 | 81 | 64 | 295 | 46 | This study |
| Cpom | 136* | 30* | 73* | 47* | 30* | 65* | 39* | 50* | 85* | 43 | Wan et al. (2019) |
| Dple | 107 | 35 | 73 | 76 | 47 | 74 | 62 | 68 | 253 | 63 | Zhan et al. (2011) |
| Tni | 143 | 51 | 122 | 71 | 68 | 85 | 74 | 91 | 274 | 55 | Chen et al. (2019b) |
| Slit | 182 | 66 | 153 | 223 | 64 | 107 | 62 | 87 | 261 | 58 | Cheng et al. (2017) |
| Harm | 122 | 57 | 105 | 76 | 54 | 120 | 71 | 79 | 257 | 53 | Song et al. (2015) |
| Bman | 94 | 37 | 94 | 64 | 48 | 69 | 56 | 74 | 218 | 59 | Xiang et al. (2018) |
| Bmor | 156 | 51 | 149 | 108 | 50 | 85 | 57 | 93 | 243 | 56 | Xia et al. (2004) |
| Msex | 164 | 66 | 137 | 103 | 51 | 89 | 76 | 79 | 261 | 67 | Kanost et al. (2016) |
| Gmel | 137 | 44 | 75 | 72 | 58 | 95 | 81 | 64 | 295 | 55 | Lange et al. (2018) |
| Ofur | 126 | 48 | 115 | 112 | 46 | 93 | 67 | 75 | 270 | 59 | Ma et al. (2020) |

\*, data from Wan et al. (2019); the other data were manually identified in our study. Pxyl, *Plutella xylostella*; Gmol, *Grapholita molesta*; Cpom, *Cydia pomonella*; Dple, *Danaus plexippus*; Tni, *Trichoplusia ni*; Slit, *Spodoptera litura*; Harm, *Helicoverpa armigera*; Bman, *Bombyx mandarina*; Bmor, *Bombyx mori*; Msex, *Manduca sexta*; Gmel, *Galleria mellonella*; Ofur, *Ostrinia furnacalis*.

## Conclusions

We assembled a genome for the OFM, which allowed us to compare genomic changes in the evolution of Lepidoptera insects. Overall, the OFM genome provides a useful resource for understanding the genetic basis of traits that underlie the ecology of budworms and the evolutionary divergence and convergence of these insects more generally. This genomic resource may also ultimately be useful for the management of OFM, such as through the identification of novel targets for chemical control and resistance monitoring.

## Data Availability Statement

The whole genome assembly has been deposited in the Genome repository (accession numbers: CP053120-CP053147) under NCBI BioProject PRJNA627114.

## Author contributions

Shu-Jun Wei conceived and designed the study; Li-Jun Cao, Wei Song, Lei-Yue, Shao-Kun Guo, Jin-Cui Chen, Ya-Jun Gong conducted the experiments; Wei Song, Li-Jun Cao and Shu-Jun Wei analyzed the data; Shu-Jun Wei, Wei Song and Li-Jun Cao discussed the results and wrote the manuscript.

## Acknowledgments

We thank Qiang Gao for his help in the assembly of Hi-C data. This research was supported by the National Natural Science Foundation of China (31901884), Joint Laboratory of Pest Control Research Between China and Australia (Beijing Municipal Science & Technology Commission), and Beijing Key Laboratory of Environmentally Friendly Pest Management on Northern Fruits (BZ0432).

