## Supplemental Tables and Figure for "The oriental fruit moth genome provides insights into evolutionary adaptation of tortricid pests"

**Table S1** Summary statistics of generated sequence data

| Library name | Experiment title | Sequencing instrument | Total bases (bp) | Coverage |
|---|---|---|---|---|
| GM_PE | DNA Pair-End (PE) library | Illumina HiSeq X10 | 48722298600 | 94.11 |
| GM_ONT | DNA Oxford NanoPore library | Oxford NanoPore | 53706537032 | 103.74 |
| GM_HiC | DNA Hi-C library | Illumina HiSeq X10 | 120847620900 | 233.43 |
| GM_Egg | RNA-Seq library | Illumina NovaSeq | 4843131000 | 9.35 |
| GM_Larve | RNA-Seq library | Illumina NovaSeq | 4345527750 | 8.39 |
| GM_Pupa | RNA-Seq library | Illumina NovaSeq | 4517857500 | 8.73 |
| GM_Adult | RNA-Seq library | Illumina NovaSeq | 5817045150 | 11.24 |

**Table S2** Summary of non-coding RNAs in genomes of *Grapholita molesta*

| Class | Type | Number |
|---|---|---|
| rRNA count | 8s_rRNA | 61 |
| | 5s_rRNA | 63 |
| tRNA Count | tRNAs decoding Standard 20 AA | 8853 |
| | Selenocysteine tRNAs (TCA) | 14 |
| | Possible suppressor tRNAs (CTA, TTA) | 13 |
| | tRNAs with undetermined/unknown isotypes | 163 |
| | Predicted pseudogenes | 20743 |
| Total tRNAs | | 29786 |
| tRNAs with intron | | 203 |

**Table S3** Summary of repeat elements in genome of *Grapholita molesta*

| Items | No. elements | Length (bp) | Percentage of sequence (%) |
|---|---|---|---|
| **Retroelements** | 231642 | 41362559 | 7.99 |
| SINEs | 26922 | 4947739 | 0.96 |
| LINEs | 200582 | 33924345 | 6.55 |
| LTR elements | 4138 | 2490475 | 0.48 |
| **DNA transposons** | 8670 | 1730875 | 0.33 |
| Unclassified | 25907 | 3076572 | 0.59 |
| Total interspersed repeats | | 46170006 | 8.92 |
| **Small RNA** | 19652 | 4014531 | 0.78 |
| **Satellites** | 2 | 267 | 0 |
| **Simple repeats** | 135315 | 5832340 | 1.13 |
| **Low complexity** | 19218 | 906543 | 0.18 |

Note: SINEs, short interspersed nuclear elements; LINEs, long interspersed nuclear elements; LTR, long terminal repeat elements.

**Table S4** Orthofinder statistics overall among 16 species

| | |
|---|---|
| Number of genes | 339228 |
| Number of genes in orthogroups | 320445 |
| Number of unassigned genes | 18783 |
| Percentage of genes in orthogroups | 94.5 |
| Percentage of unassigned genes | 5.5 |
| Number of orthogroups | 14425 |
| Number of species-specific orthogroups | 427 |
| Number of genes in species-specific orthogroups | 1785 |
| Percentage of genes in species-specific orthogroups | 0.5 |
| Mean orthogroup size | 22.2 |
| Median orthogroup size | 18 |
| G50 (assigned genes) | 29 |
| G50 (all genes) | 28 |
| O50 (assigned genes) | 2958 |
| O50 (all genes) | 3286 |
| Number of orthogroups with all species present | 4948 |
| Number of single-copy orthogroups | 144 |

G50, the number of genes in the orthogroup such that 50% of genes are in orthogroups of that size or larger; O50, the smallest number of orthogroups such that 50% of genes are in orthogroups of that size or larger.

**Table S5** Orthofinder statistics per species

| Content | Agam | Agla | Bman | Bmor | Cpom | Dple | Dmel | Gmel | Harm | Msex | Ofur | Pxyl | Slit | Tcas | Tni | Gmol |
|---|---|---|---|---|---|---|---|---|---|---|---|---|---|---|---|---|
| Number of genes | 14102 | 20632 | 19224 | 22510 | 17184 | 19762 | 30440 | 17306 | 21035 | 21890 | 23871 | 21674 | 24319 | 22610 | 23658 | 19011 |
| Number of genes in orthogroups | 12174 | 19115 | 19078 | 22191 | 16417 | 19328 | 25273 | 16629 | 20701 | 21289 | 22912 | 20230 | 22828 | 20785 | 23237 | 18258 |
| Number of unassigned genes | 1928 | 1517 | 146 | 319 | 767 | 434 | 5167 | 677 | 334 | 601 | 959 | 1444 | 1491 | 1825 | 421 | 753 |
| Percentage of genes in orthogroups | 86.3 | 92.6 | 99.2 | 98.6 | 95.5 | 97.8 | 83 | 96.1 | 98.4 | 97.3 | 96 | 93.3 | 93.9 | 91.9 | 98.2 | 96 |
| Percentage of unassigned genes | 13.7 | 7.4 | 0.8 | 1.4 | 4.5 | 2.2 | 17 | 3.9 | 1.6 | 2.7 | 4 | 6.7 | 6.1 | 8.1 | 1.8 | 4 |
| Number of orthogroups containing species | 8256 | 9117 | 10226 | 10685 | 8883 | 10256 | 8353 | 10284 | 10666 | 10582 | 10835 | 9813 | 10951 | 9142 | 10621 | 9949 |
| Percentage of orthogroups containing species | 57.2 | 63.2 | 70.9 | 74.1 | 61.6 | 71.1 | 57.9 | 71.3 | 73.9 | 73.4 | 75.1 | 68 | 75.9 | 63.4 | 73.6 | 69 |
| Number of species-specific orthogroups | 41 | 25 | 4 | 4 | 12 | 11 | 163 | 4 | 10 | 18 | 19 | 36 | 14 | 42 | 11 | 13 |
| Number of genes in species-specific orthogroups | 199 | 93 | 11 | 10 | 37 | 51 | 631 | 10 | 27 | 71 | 66 | 194 | 47 | 189 | 32 | 117 |
| Percentage of genes in species-specific orthogroups | 1.4 | 0.5 | 0.1 | 0 | 0.2 | 0.3 | 2.1 | 0.1 | 0.1 | 0.3 | 0.3 | 0.9 | 0.2 | 0.8 | 0.1 | 0.6 |
| Gene family expansion/contractions | 1857 | 825 | 1140 | 1247 | 928 | 909 | 946 | 901 | 991 | 955 | 983 | 854 | 989 | 818 | 972 | 883 |

Note: Agam: *Anopheles gambiae*; Agla: *Anoplophora glabripennis*; Bman: *Bombyx mandarina*; Bmor: *Bombyx mori*; Cpom: *Cydia pomonella*; Dple: *Danaus plexippus*; Dmel: *Drosophila melanogaster*; Gmel: *Galleria mellonella*; Harm: *Helicoverpa armigera*; Msex: *Manduca_sexta*; Ofur: *Ostrinia furnacalis*; Pxyl: *Plutella xylostella*; Slit: *Spodoptera litura*; Tcas: *Tribolium castaneum*; Tni: *Trichoplusia ni*; Gmol: *Grapholita molesta.*

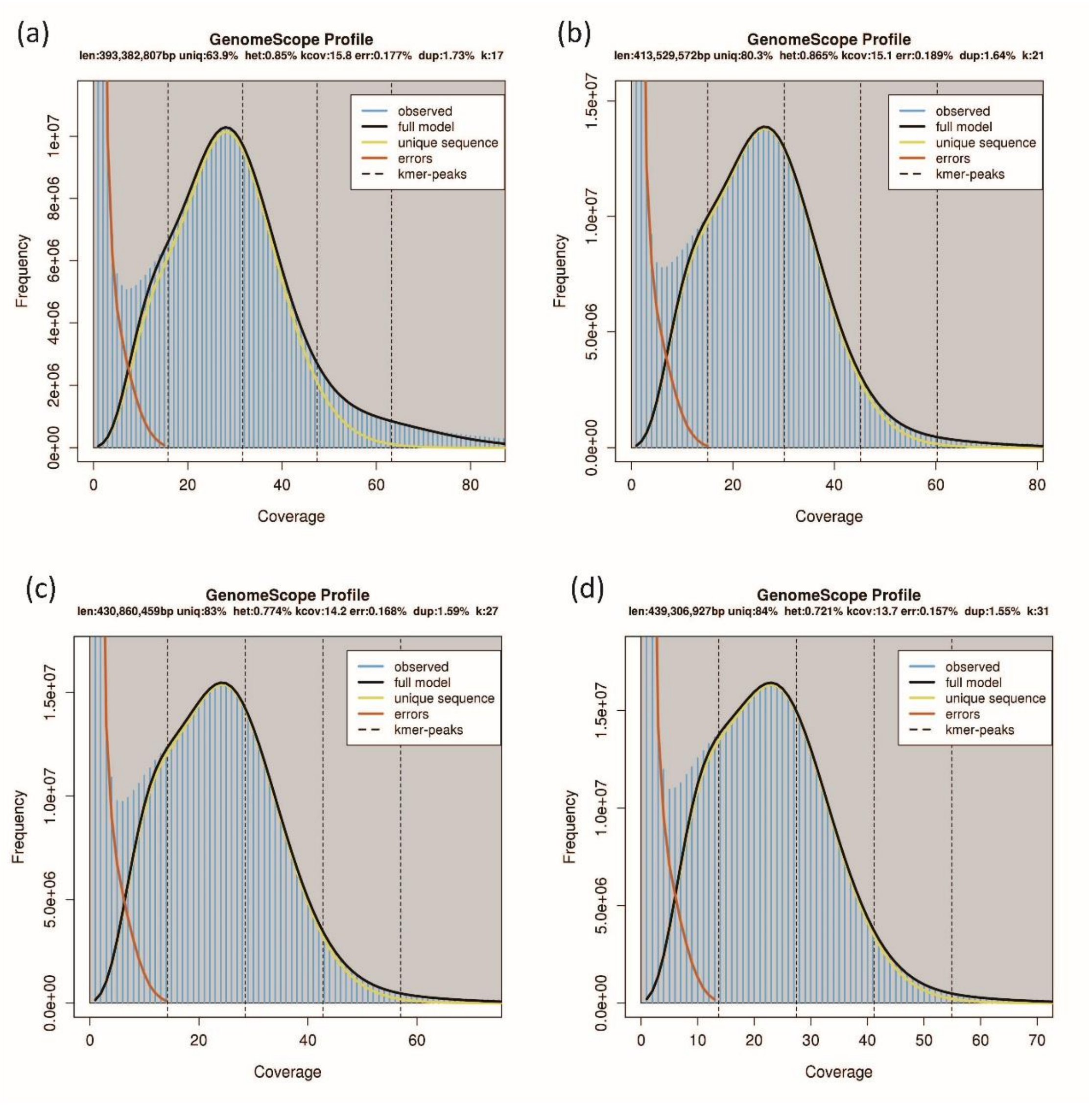


**Figure S1** Genomescope plot of kmers based on Illumina short read sequences of OFM genome when k= 17 (a), 21 (b), 27 (c) and 31 (d). Genome size (len), heterozygosity (het), error (err) and duplication (dup) were estimated and indicated in each corresponding figure.
